# Reporting and justification of sample size in translational chronic variable stress procedures: A systematic review

**DOI:** 10.1101/2024.09.26.615121

**Authors:** Crispin Jordan, Nicola Romanò, John Menzies

## Abstract

All *in vivo* studies using laboratory animals should be guided by the Three Rs: Replacement, Reduction and Refinement. The concept of Reduction is important in sample size estimation; the sample size used should allow the detection of a biologically meaningful effect size using appropriate statistical tests, but not at the expense of animal suffering. Because studies using chronic variable stress (CVS) procedures deliberately impose suffering, we reasoned that Three Rs principles would be a strong consideration in experimental design. To explore this, we conducted a systematic review of CVS studies to ask whether a biologically meaningful effect size was used to determine the sample size. Only one article in our sample of 385 reported doing this. Accordingly, it is questionable whether most of these studies align strongly with the principle of Reduction. While determining a biologically meaningful effect size is not always straightforward, we believe it is central to making biologically informed decisions about study design and interpretation, and we discuss possible ways forward.

## Introduction

The Three Rs (Replacement, Reduction and Refinement) provide a foundational framework for ethical animal research. Reduction refers to a “reduction in the numbers of animals used to obtain information of a given amount and precision” (1). A key reduction-related component of study design is selection of an appropriate sample size, i.e., using an appropriate number of animals to detect a particular effect size using a particular experimental and statistical design (2–4). If a study’s sample size is too small (and many are (5)), then small – but biologically meaningful – effect sizes may not be detected, and ‘statistically significant’ findings may actually be false positives (6,7). Because of this, a properly justified calculation of sample size is at the nexus of ethical responsibility and scientific credibility. Reporting a justified calculation demonstrates that researchers have considered what they want to measure, the magnitude of the effect they consider to be biologically meaningful and that they have used an appropriate experimental and statistical design to do this. Accordingly, prior to beginning a study, researchers would normally identify the experimental unit, determine the magnitude of effect size that is biologically meaningful in the context of their study, select an appropriate significance level (α) and statistical power, then use a power calculation to determine what sample size is needed to detect that effect size (7). While this approach has its limitations (8), it is not yet widely used in biomedical research (9).

Only a minority of studies report how Three Rs principles inform study design (10), and poor-quality study design has clear implications for animal welfare. All the animals involved in a study with a too-small sample size will have experienced some degree of suffering (even if it is considered by the relevant legislation and regulations as minor) in order to generate potentially unreliable data. This is unethical (2,11). Similarly, some of the animals involved in a study with a too-large sample size did not need to be involved in the study in order for the researchers to obtain reliable data (12). This is also unethical (2,11).

Most researchers are interested in making observations and finding interventions that are important biologically. Despite being emphasised by the National Centre for the Three Rs (13) and specified in the ARRIVE 2.0 guidelines (14), the concept of a biological meaningful effect size does not yet seem to be widely acknowledged or applied. This may partly be because of a strong (but misplaced) focus on achieving statistical significance rather than seeking biological significance (15). This may also be because a biologically meaningful effect size must be defined prior to designing the study, requiring the researcher to invest in understanding the biological system being studied, the appropriateness of the methods used, and the interventions tested. Doing this, especially if existing evidence is sparse or of poor quality, may reduce confidence in selecting a biologically meaningful effect size. Additionally, committing to test for an effect size that other researchers may not agree is biologically meaningful may inhibit a researcher from making this decision. Johnson et al (16) note that these concerns are frequently expressed as a question: how can I power my study to detect an unknown effect size? They reply that “a study should be powered to detect not the actual effect (which cannot anyway be known before collecting the data) but the smallest effect […] that, in the judgement of the researcher, is worth detecting”.

Previous work shows that few researchers report how the sample size was calculated. For example, after the introduction of a reporting checklist in a high-profile neuroscience journal, only 7% of articles reported using a power calculation to determine the sample size (17). Some studies have noted small improvements in checklist-associated reporting of sample size (18,19), but a similarly low prevalence of reporting of sample size calculation is documented in systematic reviews of a range of animal studies (10,20–34) and in an online ‘living review’ of transgenic animal models of Alzheimer’s disease (35).

We decided to explore the prevalence of sample size calculations in studies where deliberate imposition of suffering is an intrinsic part of the study. We reasoned that taking an ethical approach should be of even higher importance in this context, and that researchers would, therefore, be motivated to ensure their statistical and experimental design aligned with the key ethical principle of reduction. To do this, we carried out a systematic review to explore whether translational research using chronic variable stress (CVS) procedures (also called chronic unpredictable stress or chronic mild stress (36)) used a biologically meaningful effect size to calculate the sample size.

## Materials and methods

We searched PubMed for the terms “chronic variable stress” and “chronic unpredictable stress” in the Title/Abstract field. The search was done on 31st October 2023 and included all articles published from November 2018, i.e., a five-year period. Details on inclusion criteria are shown in Figure 1.

**Figure 1:**
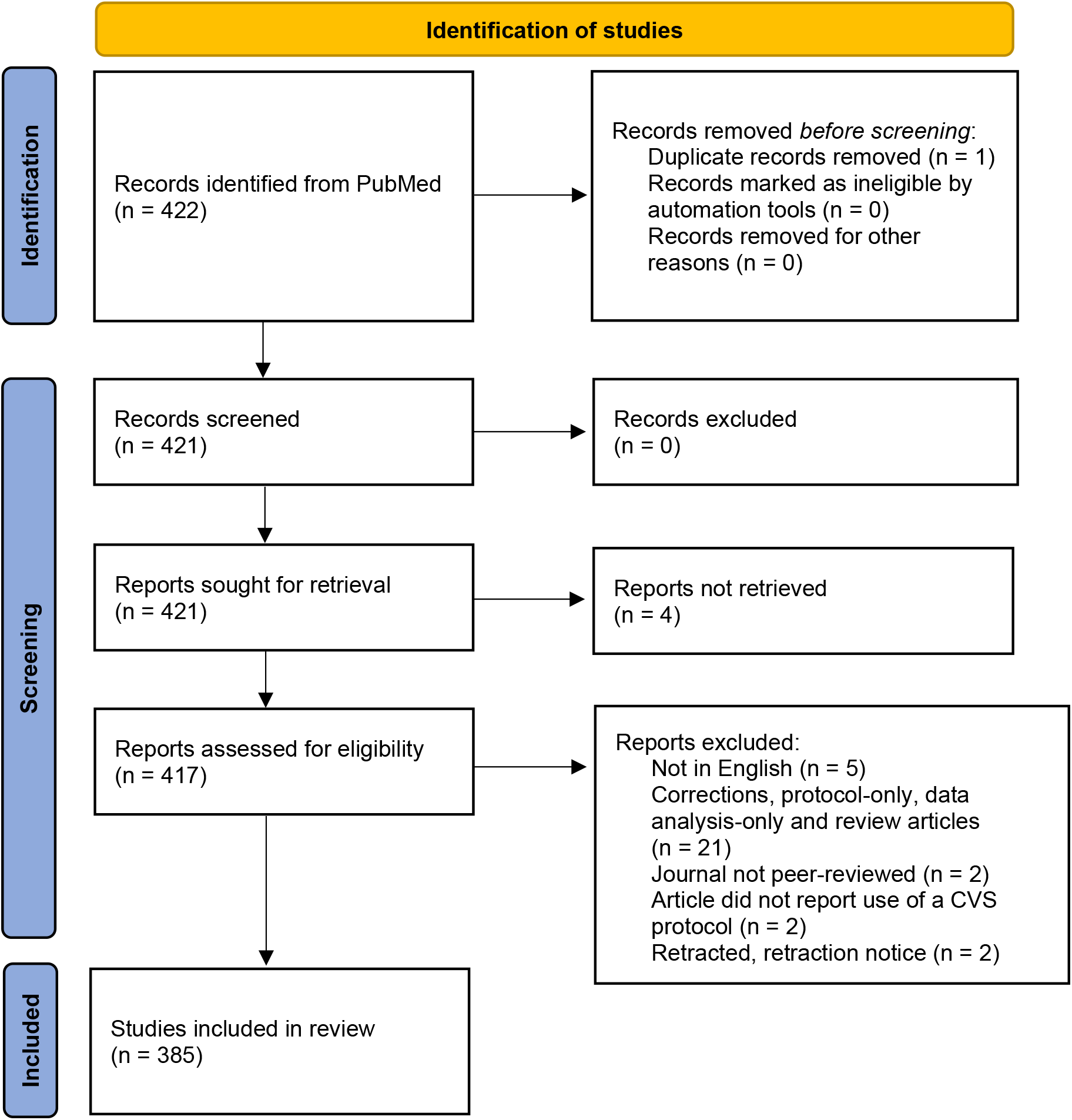
Identification of studies via databases. Based on the PRISMA 2020 Statement (37).

To explore whether and how sample size had been calculated and whether effect size was a factor in this calculation, we searched the online text of each article at the journal website and all associated supplementary files for all instances of the words: “sample”, “effect”, “group”, “size” and “power”. Journal websites were accessed between 31st October 2023 and 22nd April 2024. A single researcher recorded each instance and documented the relevant text from the article. All comments on sample size and effect size were subject to thematic analysis (38) to determine whether the full-text article contained a justification for the number of experimental units used in the experimental groups, whether this justification was based on a power calculation, and if it was, what was the source of the effect size used in that power calculation.

## Results

359 of the 385 articles (93%) included in our review reported on studies with commonly-used laboratory rodents: 199 articles (52%) used mice, 160 articles (42%) used rats, and five articles used both rats and mice. Sixteen articles used zebrafish, two used *Drosophila melanogaster*, one used California deermice (*Peromyscus californicus*), one used Japanese quail (*Coturnix japonica*) and one used cynomolgus monkeys (*Macaca fascicularis*). 180 articles (47%) did not give the species name in the title, and 13 articles (3%) did not give the species name in either the title or the abstract. Only one article in our sample (39) – describing a protocol to induce stress in *Drosophila melanogaster* – was preregistered (40).

Only one article (0.3%) reported using a biologically meaningful effect size in order to determine a sample size (41) (Figure 2; Supplementary Data 1). The corresponding author confirmed that the sample size calculation was based on a biologically meaningful effect size because of an ethical concern to use a scientifically appropriate number of rats (Prof Özlem Özmen, personal communication, 19th November 2023). Of the remaining 384 articles, 298 articles (77%) did not comment on either the sample size used and/or effect size. Of the articles that did mention sample size or effect size, twenty-one articles (6% of the total) stated sample size was based in data in previous literature or the authors’ own previous work but only one of these articles specified how that previous work informed the determination of sample size.

**Figure 2:**
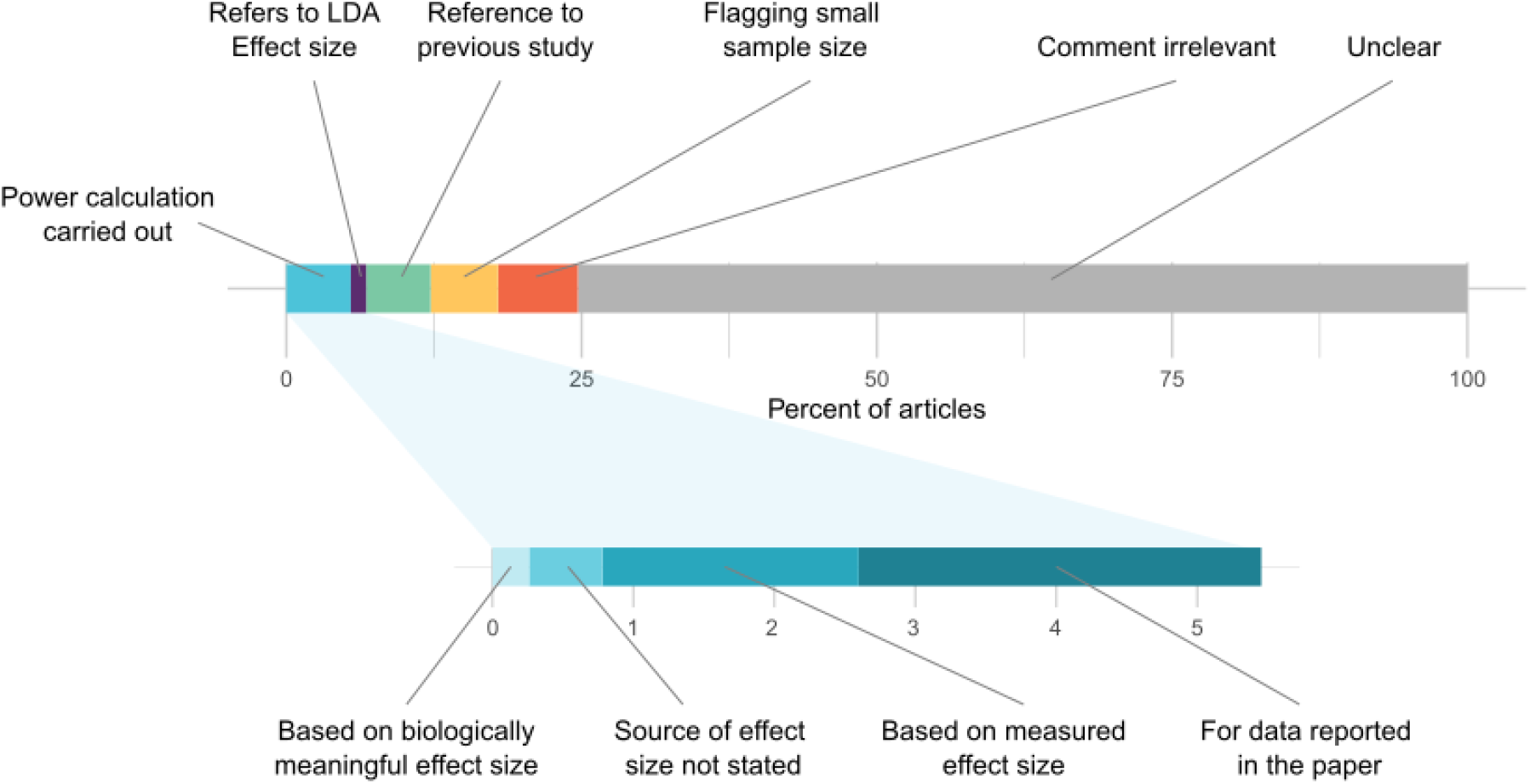
Quantitative summary of thematic analysis of sample size reporting in included articles.

Seven articles (2%) stated sample size was calculated with a power calculation using data from a previous study. Two articles stated a power calculation was used to determine the sample size but did not state the source of the effect size used. Two articles reported using Mead’s resource equation (42) to calculate the sample size.

Comments in 26 articles (7%) were not directly relevant to the calculation of the sample size (for example: “all efforts were made to reduce sample size and minimize animal suffering”). Comments made in 22 articles (6%) were to flag that the sample size used in the study may have been too small. Eleven articles (3%) carried out a post hoc power analysis, though the purpose of this was unclear as it is generally accepted that, if the p value is reported, calculating the power does not provide additional useful information (43). Five articles contained the phrase “effect size” in the context of linear discriminant analysis of effect size (LEfSe (44)).

## Discussion

Only one article of the 385 included in our review reported using a biologically meaningful effect size to calculate a sample size, but the authors did not state why the effect size was chosen. Only a further seven articles described using a power calculation to justify the sample size used. Our observations are in line with other analyses of *in vivo* animal studies where a median of only 2% of articles state how sample size was calculated (20–23,25–31,10,32,33,35). To be clear, we do not insist that a power calculation must always be used to determine sample size. For example, power analysis becomes less crucial if researchers focus their interpretation of results upon estimates of effect size and its associated uncertainty (e.g., 95% confidence intervals (CI) for the effect size) rather than upon p-values, in particular whether the p-value is less than 0.05 (45). Even then, however, methods of power analysis exist to design experiments to estimate effect size with desirably small 95% CI’s, e.g. via using simulations and replacing p< 0.05 with a desirably small 95% CI as the criterion for a ‘successful’ experiment (46). We did not calculate post hoc whether the samples sizes used meant the study designs were appropriately powered in terms of detecting the observed effect size, but in a related study (47), we note that the median sample size used in behavioural tests evaluating the effects of CVS was 10 animals per group, and that this is in line with what has been described as an ‘intuition’ of adequate sample size (48). It is not clear whether this intuition is based predominantly on statistical or biological understanding, or on practicality and/or tradition, but research has shown that published studies are frequently underpowered (5), indicating that this intuition is sometimes incorrect. At any rate, the use of incorrectly overestimated effect sizes to calculate a sample size, combined with a publication bias for ‘statistically significant’ results may lead to a biased view of what a suitable effect size is and inflate the frequency of published false positive results (49).

The apparently arbitrary selection of sample size also relates to important ethical questions. In a meta-analysis of statistical power in neuroscience research, Button et al (5) noted that average sample sizes used in a common behavioural test were capable of detecting only a relatively large effect size (Cohen’s d = 1.2), but that the likely true, smaller effect size (Cohen’s d = 0.5) needed a sample size of over 200 animals to achieve 95% power. This is a high level of statistical power, but the underlying point around ethical issues associated with small samples and small effect sizes is important; these authors noting that the total number of animals actually used across *all* the studies included in their meta-analysis was more than double the number needed to detect that small effect size in a single study. Clearly, the cumulative use of animals across a series of unreliably powered studies may use more animals than a single, well-designed study.

Significant efforts have been made to improve experimental and statistical design of translational studies, particularly the 2020 ARRIVE guidelines (14), and the adoption of reporting checklists by high-profile journals (for example, by Nature in 2013 (50)). These efforts have had a positive impact in many important regards, but less in others. A study on the impact of requiring authors to comply with ARRIVE guidelines showed a limited effect on actual compliance (19), and calls to improve the reporting of sample size calculations continue (51–54). However, if we are to encourage biological meaningfulness as a basis for this, we are faced with a challenge. When writing of using biological meaningfulness, Johnson et al suggest that studies should be designed to detect an effect that “in the judgement of the researcher, is worth detecting” (16). But how should a researcher decide what is worth detecting?

We believe most researchers want their work to matter; to matter both in the sense of having a positive impact with research and public/patient communities, and in the sense of mattering biologically and/or clinically. We argue that almost any result is of limited utility (regardless of whether it is ‘statistically significant’ (45)) if we do not have a sense of what a biologically meaningful effect size is in that context. By definition, a biologically meaningful effect size is based on understanding of biology, not based on an estimate of the ‘true’ effect size from a pilot. Accordingly, using pilot results to estimate an effect size to be detected in a ‘confirmatory’ study does not tell us how biologically meaningful this effect size is (and pilot studies using a small sample size will poorly estimate effect size). However, if exploratory work helps clarify understanding of the underlying biology, then this might help a researcher select a biologically meaningful effect size.

However, there are potential issues around using a particular sample size designed to detect a particular biologically meaningful effect size. For example, if a well-designed study uses a small sample size to detect a large biologically meaningful effect size, other researchers may dismiss this ‘low’ sample size as potentially lacking external validity. However, if researchers present a well-reasoned argument for using a relatively small sample size – one which is selected using an unbiased method and is sufficient to detect a large (and biologically meaningful) effect size – this may alleviate concerns.

So far, we have focussed on the need to identify and use the *smallest* effect size of biological interest when conducting a power analysis. However, knowledge of the smallest biologically meaningful effect is also crucial to interpret results. For example, a small p-value for an experimental treatment provides a researcher with one line of evidence that the treatment has an effect. However, if the researcher is unsure how large this effect must be to be biologically relevant, they do not have a way to judge whether their results are meaningful. For example, in a clinical trial exploring the impact of exercise on physical and mental health in patients with coronary artery disease (55), the authors identified ‘statistically significant’ effects of treatments on blood pressure, but concluded that the changes were unlikely to be clinically meaningful. In our view, this was a reasonable conclusion. Overall, regardless of a p-value obtained, meaningful interpretation can only occur with a sense of a biologically important minimal effect size (15,45). Importantly, the exact value of a biologically meaningful effect size may not be fixed. For example, new insights might change our view of what a biologically relevant effect size is, or the same effect size might be biologically relevant in one context and less irrelevant in another (yet the actual data may have the same level of statistically significance in both cases).

But how can we determine a biologically meaningful effect size? This ability arises from an understanding of the biological system, which normally derives from good-quality data scrutinised and interpreted by experts in the field. For instance, the clinical trial described above benefited from independent research that specifically aimed to determine minimally important effect sizes (56). When such data does not exist, research to determine important effect sizes becomes essential, and may include theoretical studies. One way forward is to use mathematical models of biological processes (57). Many areas of biology, like ecology and evolution, have rich and ongoing histories of using mathematical models to understand processes. Consider an example in the field of evolutionary genetics, where a researcher might wish to study a species’ potential to adapt to a local environment. Mathematical models predict that a population of a given species may evolve adaptations to a local habitat or region at the expense of being maladapted to another region or habitat when the strength of natural selection is greater than the rate of immigration between the regions (58,59). Therefore, if the researcher has estimates of immigration rates, they can use these estimates as a minimal effect size to design experiments on natural selection. Mathematical models could also be used to predict the effect size of the proposed intervention on a specific output of a biological system in the biomedical context, including behaviours (60) and physiological processes (61).

Our data demonstrate that few CVS studies use a power analysis to determine sample sizes, and even fewer use biologically informed effect sizes in their sample size calculations. This is unsatisfactory from both a scientific and ethical point of view. Although reduction may not be appropriate when it increases the level of individual suffering for the animals taking part in the experiment (62), in line with current advice (13,14), we support greater rigour in this aspect of experimental design. We further argue that meaningful interpretation of any result (regardless of p-value obtained) is impossible without a sense of what a biologically meaningful effect size is likely to be. Accordingly, if there is little sense of what a biologically meaningful effect size is, research to articulate this should be a priority.

## Supporting information

Supplementary Data 01 - thematic analysis of sample size reporting

Supplementary Data 02 - Incl-excl criteria, randomisation, blinding

## Acknowledgements

This work was supported by the Medical Research Council [grant number MR/V012290/1]. CJ thanks Prof Sally Otto (University of British Columbia) for insights into using theory to determine effect sizes. The authors acknowledge that, in our own published animal studies, we did not always practice what we preach with respect to justifying sample sizes. For the purpose of open access, the author has applied a Creative Commons Attribution (CC BY) licence to any Author Accepted Manuscript version arising from this submission.

