## Supplementary Text for "Reporting and justification of sample size in translational chronic variable stress procedures: A systematic review"

*1. Key indicators of study quality*

In addition to documenting reporting and justification of sample sizes in our sample of articles using CVS, we also systematically documented other key indicators of study quality: use of exclusion criteria, use of random allocation to experimental groups, and experimenter blinding to groups (1). To do this, the online text of each article was searched using the terms "exclu", "inclu", "criteri", "random" and "blind".

*1.1 Exclusion criteria*

Seventy-four articles (19%) reported using exclusion criteria for their study. Twenty-three articles reported criteria applied prior to data collection (for example, screening animals for their "baseline" sucrose preference (2) or behaviour in the open field test (3)), and forty-nine articles reported criteria applied after data collection (for example, excluding animals who climbed their own tails in an effort to escape from the tail suspension test (4)). Two articles listed both types of exclusion criteria. In line with other systematic reviews on translational studies, we found that only 19% of articles reported exclusion criteria for their study (5,6) (Supplementary Data 2). Twenty-five articles (6%) described excluding data points that were deemed to be outliers, usually defined as being more than two standard deviations away from the mean. We note that data points should not be excluded just because their value is distant from the mean. This is because deliberately reducing the variability in one's data by removing certain data points can disfavour the null hypothesis and increase the risk of false positives (7). Moreover, excluding data based on *a priori* expectations of what data 'should' look like undermines efforts to achieve unbiased understanding of a process; the world is full of unusual phenotypes that reflect biological reality, and removing unexpected (but otherwise reliable) data distorts understanding. If unusual data points violate a statistical test's assumptions, other forms of analysis are often available (for example, bootstrapping (8)).

*1.2 Randomisation*

178 articles (46%) reported that experimental units were allocated randomly to the control and CVS groups. This is higher than the median of 33% reported in other reviews (2,5,6,9–17), but still low overall. A further seventeen articles (4%) reported random allocation to groups *after* the CVS procedure had taken place (Supplementary Data 2).

*1.3 Blinding*

133 articles (35%) reported that experimenters were blinded during the collection and/or analysis of data. This is somewhat higher than the median of 21% reported in other reviews (2,5,6,9–20). However, in twenty-one of those articles, experimenters were not blinded during at least one of the behavioural tests used to evaluate the effects of CVS. Four articles (1%) reported that experimenters were not blinded throughout. In the remaining 64% of articles, it was unclear whether blinding was used (Supplementary Data 2).

*2. Limitations*

Our study has a number of limitations. Firstly, we did not preregister our protocol design. Secondly, by setting strict inclusion criteria we may not capture all relevant articles. Setting these criteria is intended to reduce selection bias, but translational procedures that impose stressors over long periods are known by a variety of names (21), not just by our search terms, so the included articles represent a sample of all relevant studies published in that time frame. Decisions on inclusion/exclusion, data collection and thematic analyses were done manually by a single researcher, this may introduce biases (22,23).
